# Quantitative reconstruction of time-varying 3D cell forces with traction force optical coherence microscopy

**DOI:** 10.1101/444240

**Authors:** Jeffrey A. Mulligan, Xinzeng Feng, Steven G. Adie

## Abstract

Cellular traction forces (CTFs) play an integral role in both physiological processes and disease, and are a topic of interest in mechanobiology. Traction force microscopy (TFM) is a family of methods used to quantify CTFs in a variety of settings. State-of-the-art 3D TFM methods typically rely on confocal fluorescence microscopy, which can impose limitations on acquisition speed, volumetric coverage, and temporal sampling or coverage. In this report, we present the first quantitative implementation of a new TFM technique: traction force optical coherence microscopy (TF-OCM). TF-OCM leverages the capabilities of optical coherence microscopy and computational adaptive optics (CAO) to enable the quantitative reconstruction of 3D CTFs in scattering media with minute-scale temporal resolution. We applied TF-OCM to quantify CTFs exerted by isolated NIH-3T3 fibroblasts embedded in Matrigel, with five-minute temporal sampling, using images which spanned a 500×500×500 μm^3^ field-of-view. Due to the reliance of TF-OCM on computational imaging methods, we have provided extensive discussion of the underlying equations, assumptions, and failure modes of these methods. TF-OCM has the potential to advance studies of biomechanical behavior in scattering media, and may be especially well-suited to the study of cell collectives such as spheroids, a prevalent model in mechanobiology research.

## Introduction

The field of mechanobiology seeks to understand the role of mechanical interactions and forces in both physiological processes and disease. Cellular traction forces (CTFs) are a topic of particular interest to mechanobiology researchers, as CTFs have been shown to play an integral role in many settings, including metastasis^1^, angiogenesis^2,3^, and collective cell migration^4,5^. As a result, several tools and techniques have been developed to enable the measurement of CTFs under a variety of conditions and settings^6-9^. Traction force microscopy (TFM) comprises a diverse family of methods used to quantify CTFs, based upon the optical measurement of CTF-induced deformations in the surrounding environment. TFM has enabled the discovery of several important biological findings, such as the association of strong CTF generation with the metastatic potential of cancer cells^1^, and the finding that growth cones of neurons from the peripheral nervous system exert significantly stronger forces than those of neurons from the central nervous system^10^.

As both mechanobiology and TFM continue to develop, there is a growing demand for the application of TFM in settings that pose technical challenges for the imaging systems that TFM relies upon. A growing prevalence of TFM performed in 3D environments (and the associated need for 3D imaging) has been motivated by the fact that cell behavior can vary greatly between 2D and 3D environments^11-14^. As the mechanical behavior of single cells and cell collectives span a wide range of spatiotemporal scales (i.e., micrometers to millimeters, and minutes to days)^15-18^, there is a need for high resolution imaging over large (volumetric) fields-of-view that can be achieved with high temporal sampling and/or repetition over extended durations. Finally, there is a growing interest in measuring CTFs within biopolymer substrates (e.g., collagen and fibrin)^2, 19-21^ that can introduce additional complications for imaging (e.g., optical scattering), as well as the characterization of mechanical properties and CTF reconstruction^9^. Confocal fluorescence microscopy is the dominant imaging modality for performing 3D TFM. However the limitations of this modality may restrict the range of experimental conditions in which 3D TFM is feasible. These obstacles include a limited penetration depth (of a few hundred micrometers) in scattering media, long measurement times for the acquisition of large volumes, and complications posed by photobleaching and phototoxicity.

Motivated by the developing needs of the TFM field and by the limitations of current technologies, we previously proposed a TFM method based upon optical coherence microscopy (OCM)^15^, a variant of optical coherence tomography (OCT) with high transverse resolution. The method, which we named traction force optical coherence microscopy (TF-OCM)^15^, leverages multiple advantages offered by OCM and computed imaging methods to enable the quantitative reconstruction of 3D CTFs with high temporal resolution in scattering media. These advantages include a rapid (minute-scale) volumetric acquisition rate provided by Fourier domain OCM systems, focal plane resolution over an extended depth-of-field achieved with the aid of computational adaptive optics (CAO)^22^, and label-free imaging at near-IR wavelengths to mitigate scattering and photobleaching/phototoxicity concerns. In Ref. [15], we demonstrated the feasibility of TF-OCM by showing that OCM images reconstructed with CAO procedures could be used to measure time-varying deformations induced by CTFs exerted within a 3D hydrogel substrate. However, we had not yet developed the full experimental and data processing methods required to obtain quantitative reconstructions of time-varying CTFs with TF-OCM.

In this study, we expand upon the methods reported in Ref. [15], and present a complete quantitative implementation of TF-OCM. With this new technique, we quantified time-varying 3D CTFs exerted by isolated NIH-3T3 fibroblasts embedded within a Matrigel substrate. This was achieved by analyzing OCM images acquired with five-minute temporal sampling, spanning a 500×500×500 μm^3^ field-of-view (FOV). In order to obtain OCM images suitable for TFM, we developed a computational image formation procedure which mitigates a variety of detrimental image artifacts. Although this procedure was developed for TF-OCM, the underlying mechanisms and techniques are relevant to other applications of computed OCT/OCM imaging which require geometrically accurate reconstructions of object structure. We have therefore provided extensive discussion of the equations, assumptions, and failure modes of these methods. High-level results and discussions regarding TF-OCM and computed imaging appear in the main text below, while detailed equations and discussions may be found in the Supplementary Methods and Supplementary Discussion sections of this report. The results shown here mark the first quantitative implementation of TF-OCM, and set the stage for future work toward the application of TF-OCM to more complex model systems, such as multicellular spheroids.

## Overview of TF-OCM

Detailed descriptions of our methods are available in the Methods and Supplementary Methods sections of this report. However, a brief overview of TF-OCM here is pertinent to interpreting the results in the sections that follow.

TFM does not measure CTFs directly. Instead, CTFs are numerically reconstructed based upon how they deform the surrounding environment. CTF reconstruction requires the measurement of three key pieces of information: 1) substrate mechanical properties, 2) substrate deformations in response to CTFs, and 3) boundary conditions (e.g., environment boundaries and cell geometry). The first is obtained through mechanical characterization of the substrate material (e.g., polyacrylamide, Matrigel, collagen, etc.). The latter two are obtained through a combination of optical imaging and appropriate assumptions/constraints. To measure substrate deformations in particular, the sample structure must be known from both a ‘deformed state’ (when CTFs are present) and a ‘reference state’ (when CTFs are not present). The ‘reference state’ is typically induced by a chemically-mediated inhibition of CTFs at the end of an experiment. Once the three key pieces of information are known, numerical methods are used to reconstruct the CTFs. This entire procedure can be performed in a variety of ways, creating opportunities for the development of new methods, such as the TF-OCM technique presented here^9,15^.

Our implementation of TF-OCM follows these same general procedures. Here, NIH-3T3 fibroblasts were embedded in a Matrigel susbtrate containing scattering polystyrene beads. The cells exerted contractile CTFs, causing the Matrigel to deform, resulting in displacement of the scattering beads from their resting (i.e., reference) positions. A contractility inhibitor (cytochalasin D) was then added to the culture. In response, the cells relaxed, allowing the scattering beads to return to their resting positions (due to the elasticity of the substrate). This process was observed over a three-hour period with five-minute temporal sampling using a spectral domain OCM imaging system. Volumetric images (spanning a 500×500×500 μm^3^ FOV) were reconstructed using a customized procedure which combined CAO and additional computational methods^22-25^. These images were then used to obtain measurements of time-varying bead displacements, and to generate time-varying discrete meshes of cell surfaces. These data, in addition to the mechanical properties of the Matrigel substrate (as characterized by bulk rheology), were fed into a previously reported finite element method (FEM) procedure^26^ to obtain quantitative reconstructions of time-varying 3D CTFs.

The computational workflow underlying TF-OCM is a union of two modules, which are depicted by the flowchart in Fig. 1. The computational image formation module generates a time series of high resolution, volumetric OCM images of a cell and its surroundings. The TFM module uses this image data to quantify time-varying 3D CTFs. It is important to note that, although CAO can restore focal plane resolution throughout an imaged volume, the structure of the resulting image is not necessarily a reliable representation of the true sample structure. This can be detrimental to the accurate measurement of substrate deformations and subsequent quantification of CTFs. The numerous steps preceding CAO were used to mitigate factors that corrupt the image formation process, and thus played a critical role in our implementation of TF-OCM. The results associated with Figs. 2 and 3 show the impact of a selection of these procedures and have relevance to the field of computed OCT/OCM imaging. The results associated with Figs. 4-7 are specific to TF-OCM.

**Figure 1.**
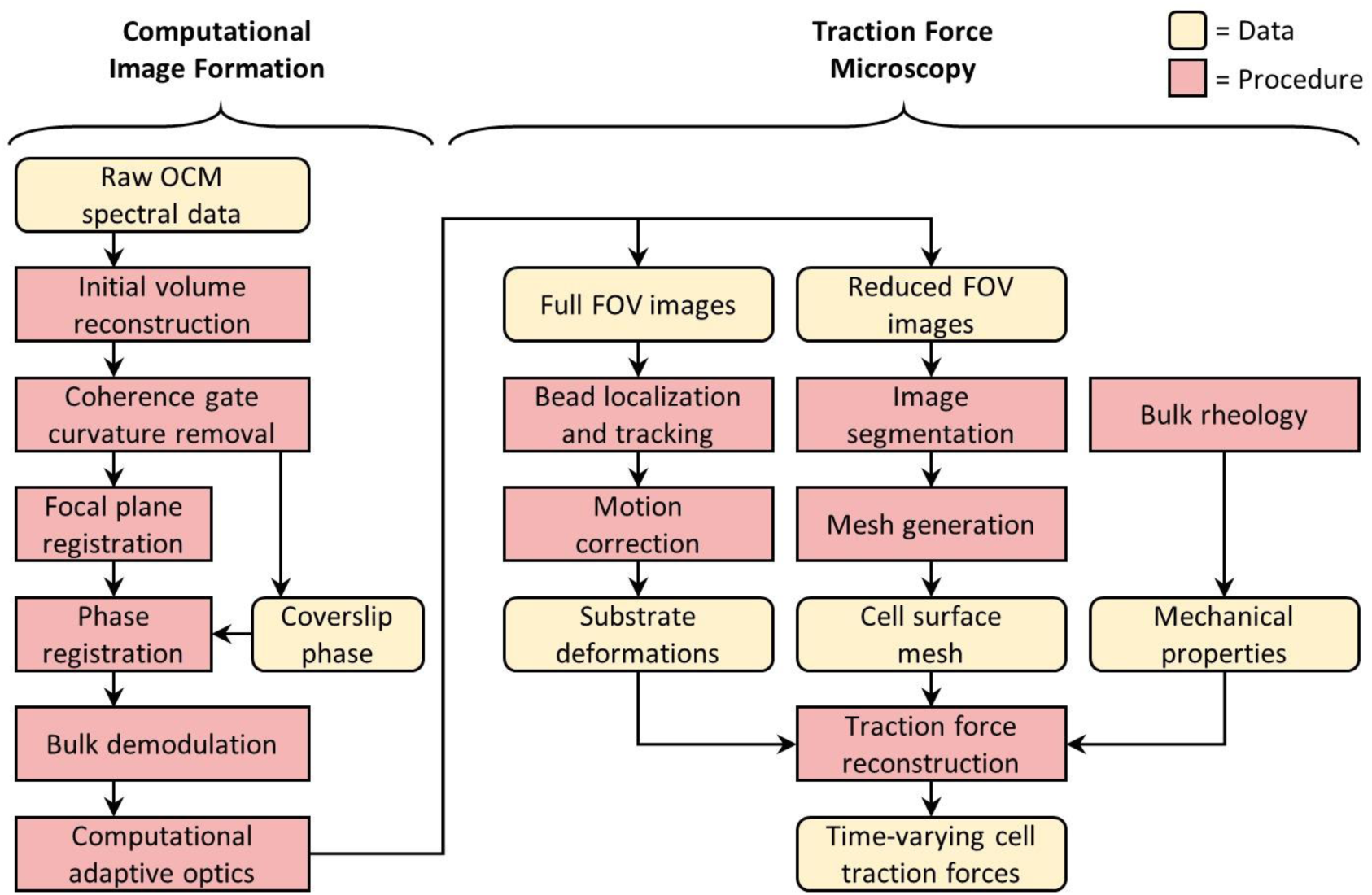
Data processing workflow of traction force optical coherence microscopy. A time series of volumetric images is reconstructed via the computational image formation module (left). These images are then used to quantify time-varying 3D CTFs via the traction force microscopy module (right).

**Figure 2.**
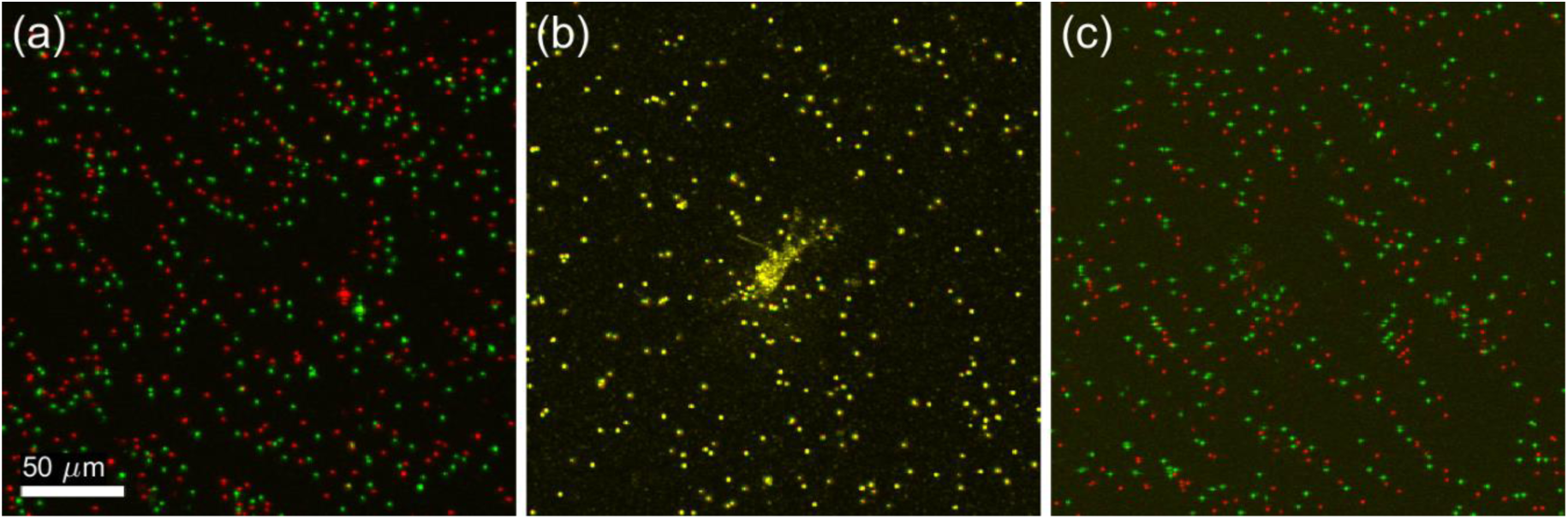
Shearing artifacts in computationally refocused images. Panels (a), (b), and (c) were obtained from 3 separate depths, located 200 μm above, at, and 200 μm below the focal plane, respectively. The red channel corresponds to images obtained using our phase registration and bulk demodulation procedures, whereas the green channel corresponds to images obtained without these procedures. The green channel exhibits a depth-dependent translation artifact, corresponding to a shearing of the reconstructed volumetric image. Additional spatial and temporal variations in this artifact are visible in Supplementary Movie 1.

**Figure 3.**
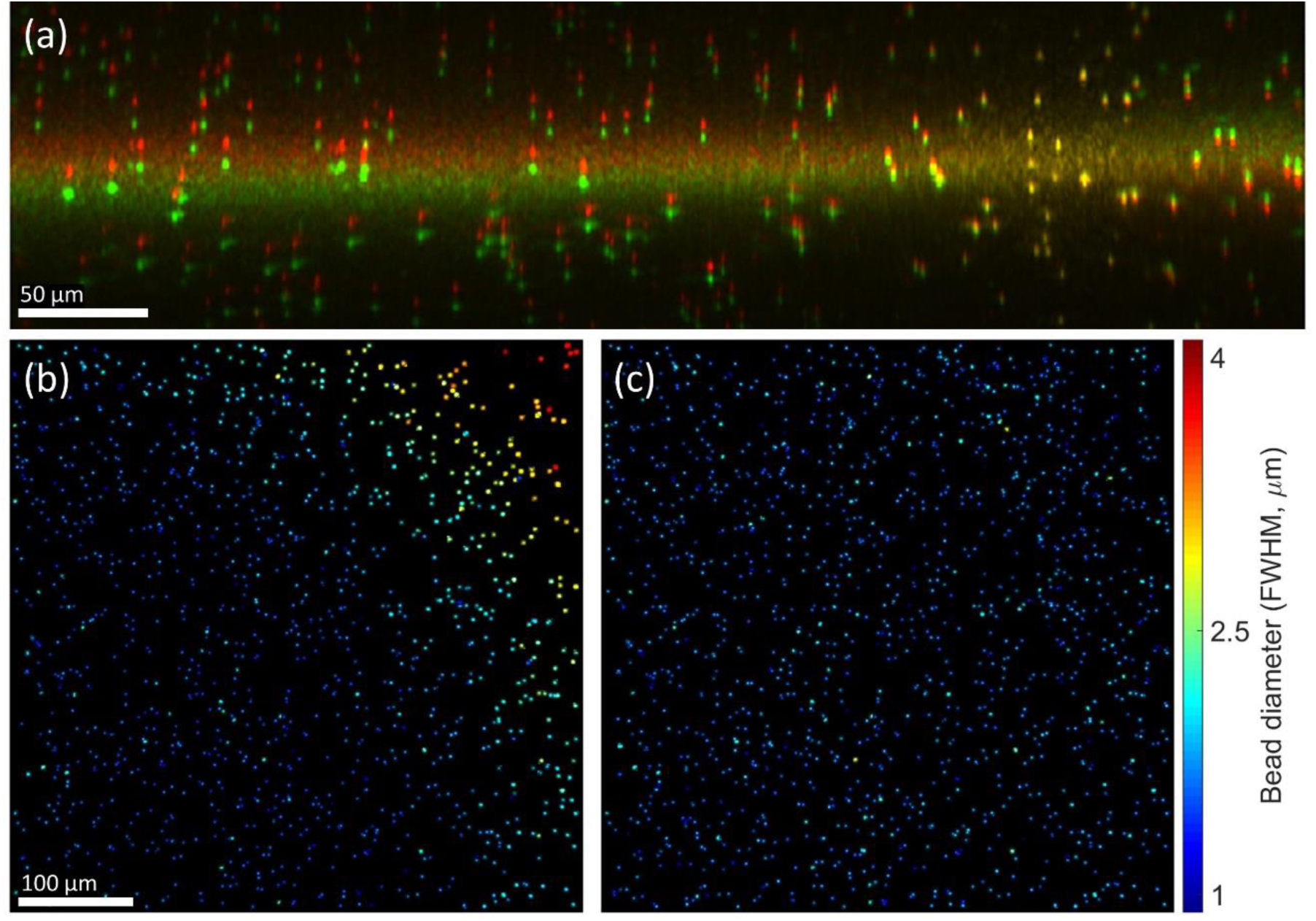
Effects of focal plane registration on volumetric image reconstructions. (a) Cross-section of a volumetric image obtained with (red channel) and without (green channel) focal plane registration. The bright horizontal band spanning both channels corresponds to the focal plane. (b,c) *En face* planes obtained from a region centered 50 μm above the focal plane in volumetric images reconstructed with CAO. Only beads which exhibited little to no overlap with neighboring beads were retained in these images. The methods for generating these panels are detailed in the Supplementary Methods. (b) *En face* plane reconstructed without focal plane registration, exhibiting a transverse resolution which varies across the FOV. (c) *En face* plane reconstructed with focal plane registration, resulting in a uniform transverse resolution. FWHM indicates the full-width-at-half-maximum diameter of the polystyrene beads shown in (b) and (c). This value is not a measurement of the post-CAO imaging resolution, but is directly correlated to resolution and is therefore used here to demonstrate the impact of focal plane registration. Color bar applies only to (b) and (c).

**Figure 4.**
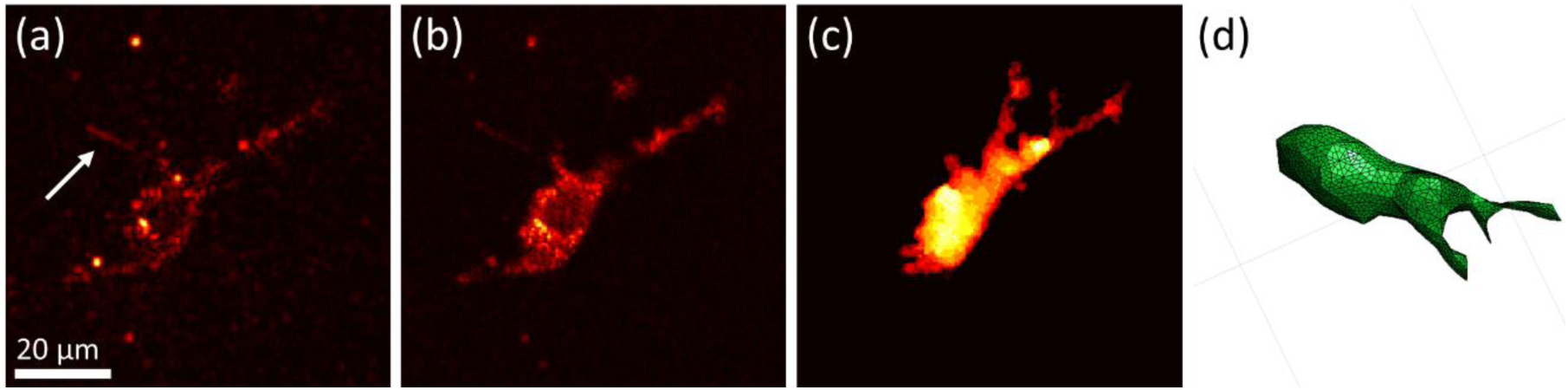
Automated segmentation of cell bodies aided by speckle reduction. (a) *En face* plane extracted from a single depth in a single volumetric image. The imaged NIH-3T3 fibroblast exhibits significant speckle artifacts, which hinder automated segmentation. Arrow indicates cellular structure not retained by our segmentation procedure (see text for details). (b) The same *en face* plane as in (a), after combining eight sequential volumetric acquisitions. Speckle contrast is reduced, allowing for segmentation via K-means clustering. (c) Summation projection of the 3D segmented volume, which approximates the cell body. (d) 3D mesh of the cell body, generated from the data depicted in (c). Note that (d) was generated from a different viewing angle as (a-c) to more clearly depict the cell’s 3D shape.

**Figure 5.**
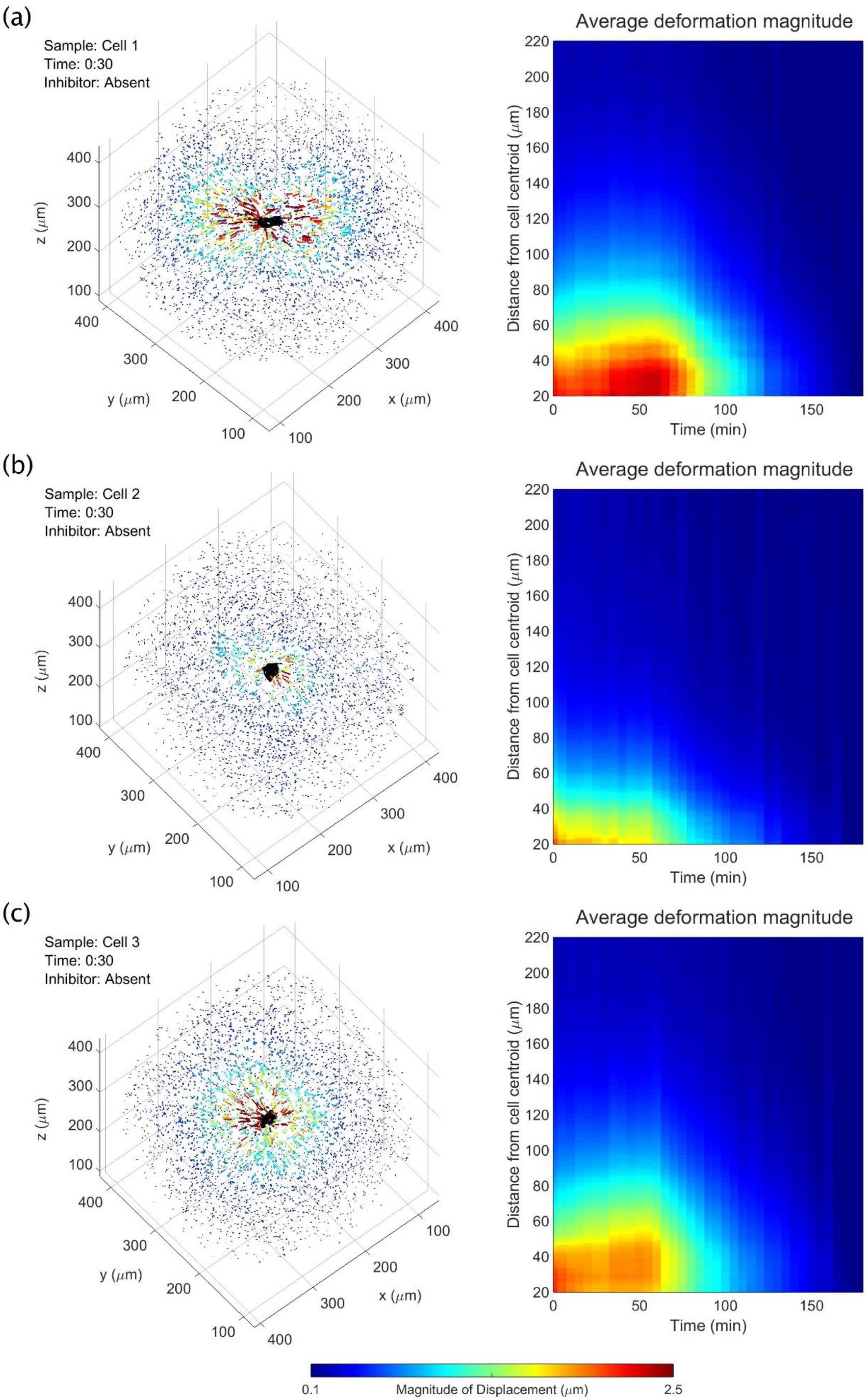
Time-varying, 3D substrate deformations measured with TF-OCM. (a,b,c) Bead displacement data for three NIH-3T3 fibroblasts. Left panels depict bead displacements at the time point immediately preceding the addition of the contractility inhibitor, cytochalasin D. Arrows indicate bead displacements with respect to their ‘reference’ positions (i.e., resting positions in the absence of CTFs). Arrow lengths are exaggerated for visualization. Animations over time may be found in Supplementary Movie 2. Right panels depict the mean magnitude of bead displacement as a function of time and distance from the cell centroid (see the Supplementary Methods for details on this calculation). The bead localization sensitivity along the *x*, *y*, and *z* axes were 37 nm, 32 nm, and 86 nm, respectively (described in the Methods section).

**Figure 6.**
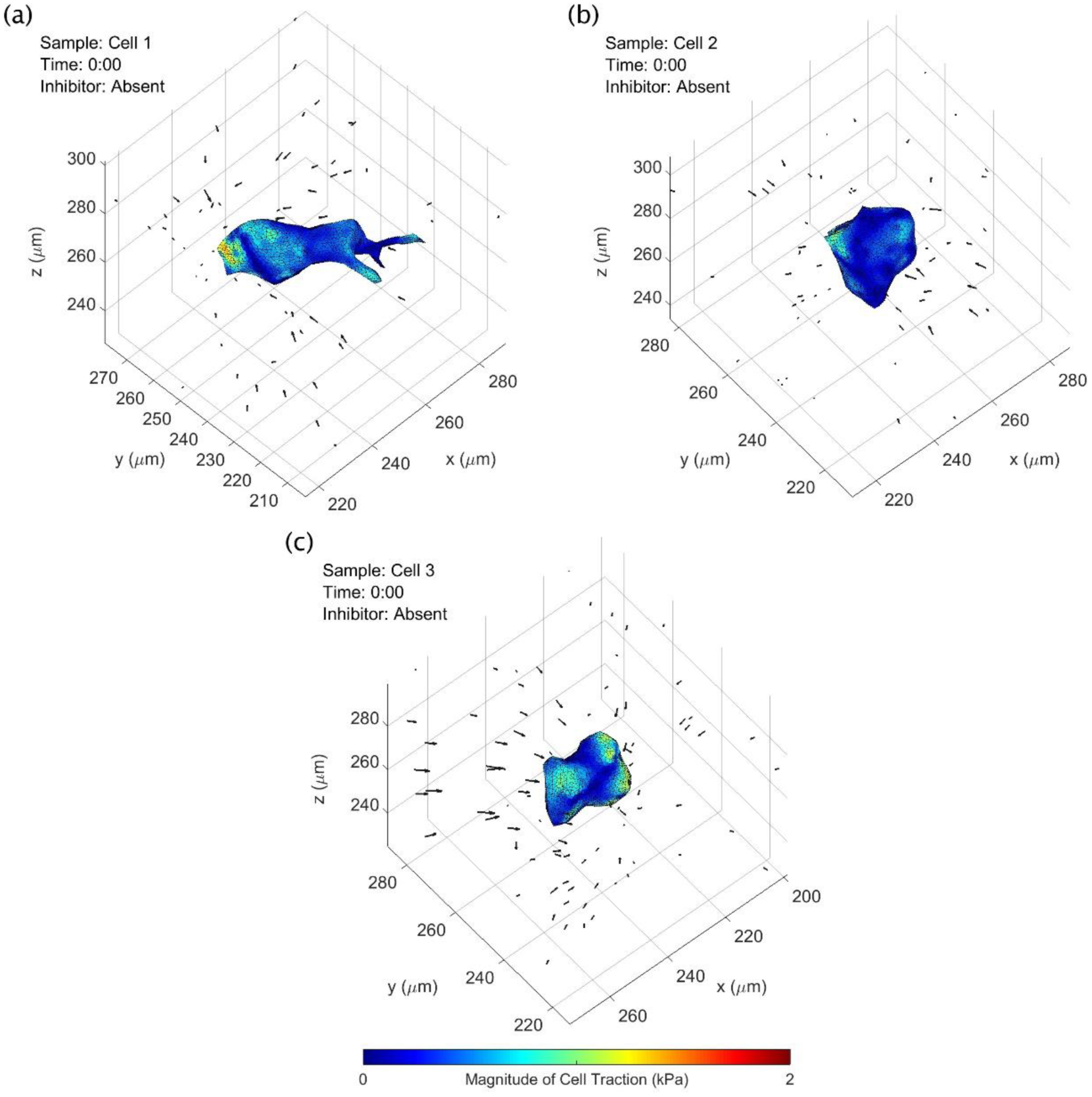
CTF reconstructions at a single time point for three different NIH-3T3 fibroblast cells. A time-lapse animation of this figure is provided in Supplementary Movie 3. Black arrows indicate measured bead displacements with respect to their reference positions. See text for details.

**Figure 7.**
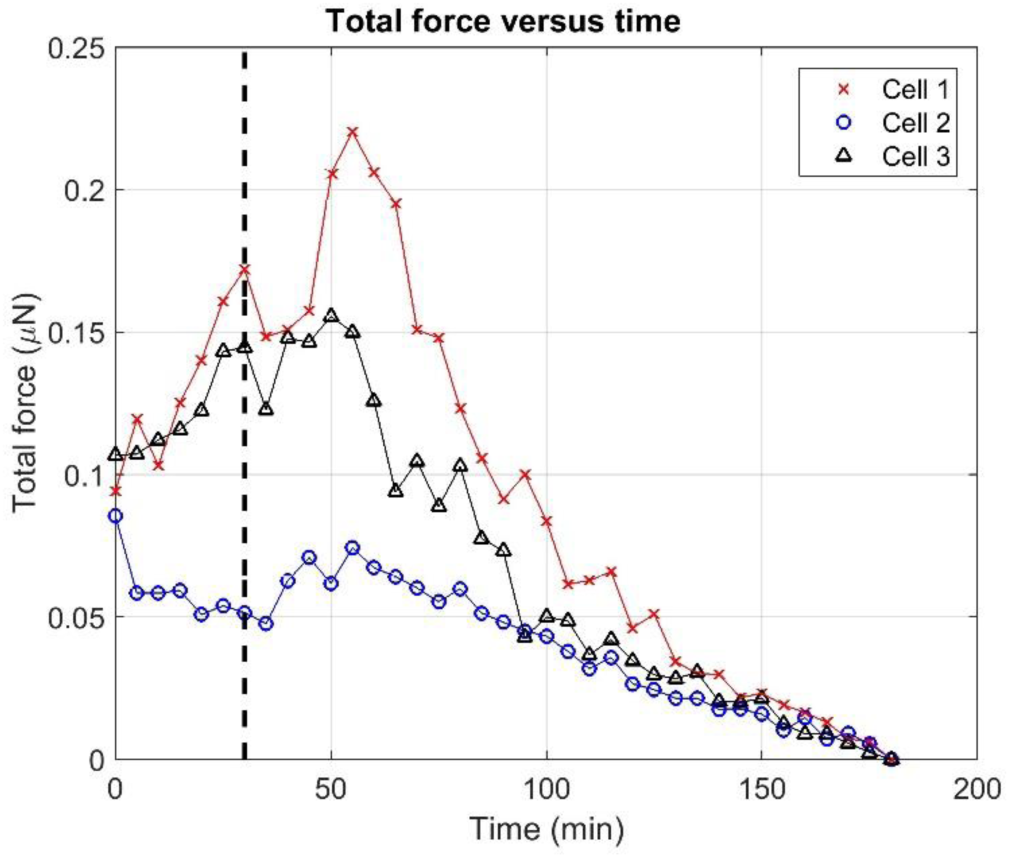
Total force exerted by NIH-3T3 fibroblasts over time. Black dashed line indicates when the contractility inhibitor (cytochalasin D) was introduced to the samples 30 minutes into the experiment. See the Supplementary Methods for a description of the measurement of ‘total force’ from 3D CTF distributions.

## Results

### Phase registration and bulk demodulation mitigate shearing and motion artifacts in computationally refocused images

As our implementation of TF-OCM relies upon the accurate reconstruction of high resolution volumes via CAO, we had to design our computational image formation module to ensure that interferometric phase instability and other factors would not corrupt the image data^23,24^. Figure 2 and Supplementary Movie 1 demonstrate the impact and necessity of our phase registration and bulk demodulation procedures (detailed in the Supplementary Methods) for mitigating such effects. In Fig. 2, the red and green channels (which correspond to images generated by our computational image formation module with and without these procedures, respectively) exhibit a depth-dependent transverse misalignment with respect to one another. The severity of this misalignment increases linearly with distance from the focal plane. In addition, the direction of the misalignment has opposite signs on opposite sides of the focal plane. The red and green channels are therefore distorted relative to one another by a shearing deformation (analogous to pushing against the side of a deck of cards). Supplementary Movie 1, which depicts a time lapse of the *en face* planes shown in Fig. 2, reveals additional features of this misalignment. In panels (a) and (c) of the animation, the embedded scattering particles are at depths far away from the imaged NIH-3T3 fibroblast cell, and are expected to exhibit little to no motion in the animation. This expected behavior is exhibited by the red (phase-registered and demodulated) channel. In contrast, bead positions in the green (uncorrected) channel fluctuate over time. These fluctuations are non-uniform across the FOV and are most prevalent along the left-right axis, corresponding to the slow scanning axis of our imaging system (which is known to be more susceptible to interferometric phase fluctuations ^23,24^). Given that the red channel of Supplementary Movie 1 exhibits the behavior we expect to observe (i.e., little to no motion of beads far from the cell), we conclude that our phase registration and bulk demodulation procedures are effective at mitigating shearing and motion artifacts in computationally refocused OCM images, and are therefore beneficial to the accuracy of TF-OCM. Further theoretical justification of this conclusion may be found in the Supplementary Discussion.

### Focal plane registration facilitates CAO over a wide field-of-view for systems with non-ideal optics

Non-idealities in the sample arm optics can negatively impact the reconstructed resolution and signal-to-noise-ratio (SNR) of computationally refocused OCM images, and are therefore detrimental to both the tracking of bead displacements and identification of cell boundaries in TF-OCM. In this study, non-idealities in our sample arm optics caused the axial position of the focal plane to vary as a function of transverse position (such that the focal plane appears slightly tilted and/or curved). This resulted in an imaging point spread function (PSF) that varied across the 500×500 μm^2^ transverse FOV of our images. As our implementation of CAO (detailed in the Supplementary Methods) applies a refocusing operation that is transversely invariant, it cannot account for transverse non-uniformity in the imaging PSF.

In the absence of our focal plane registration procedure, refocused images exhibited a non-uniform transverse resolution after CAO, as demonstrated in Fig. 3(b). To mitigate this effect, we devised a focal plane registration procedure (detailed in the Supplementary Methods) which maps the image data to a coordinate system in which the axial position of the focal plane is transversely invariant. (See Fig. 3(a) for a visual example of this transformation.) Following this procedure, images refocused with CAO exhibited a transverse resolution that was uniform across the transverse FOV, as in Fig. 3(c). These results show that our focal plane registration technique allows for a standard implementation of CAO to refocus volumetric data over a wide FOV (500×500 μm^2^), even when non-idealities of the optical system cause the focal plane to be tilted or curved with respect to the axes of the image data. Of course, spatially-varying CAO reconstructions may instead be employed to address this problem at the cost of greater complexity^27,28^. However, the methods used in this study may be considered as a simple alternative to be used when the tilt/curvature of the focal plane is small.

### Speckle reduction aids automated analysis of cell geometry from OCM images

As discussed previously, the reconstruction of CTFs requires knowledge of cell shape/geometry. Cell tracing based on automated image segmentation of OCM images can be difficult, due to the presence of speckle (such as that shown in Fig. 4(a)), which is ubiquitous to coherent imaging modalities like OCT/OCM, ultrasound, radar, etc. However, the alternative prospect of performing manual cell tracing would have been prohibitive to conducting TF-OCM in time-lapse settings. For example, the three time-lapse experiments reported in this preliminary study span a total of 111 distinct time points. Performing manual 3D tracing typically would have required manual 2D tracing of at least 15 slices from the volumetric image corresponding to each time point. For this study alone, that would have entailed performing manual tracing for more than 1600 individual 2D images. To overcome this obstacle and keep TF-OCM feasible for larger experiments in the future, we employed speckle reduction methods to aid the automated analysis of cell geometry from OCM images. To reduce speckle artifacts in and around the cell body, we combined multiple sequentially-acquired OCM images (as detailed in the Methods), which resulted in images like the one shown in Fig. 4(b). The reduction in speckle provided by our technique enabled automated image segmentation and identification of the cell body via K-means clustering^29^, yielding 3D segmented regions like that depicted in Fig. 4(c). Segmented regions were converted to 3D discrete meshes, as in Fig. 4(d), for use in our FEM solver.

Note that some of the finer structures visible in Figs. 4(a) and 4(b) were not retained in the segmentation shown in Fig. 4(c). We found that our speckle reduction method can suppress cellular structures which exhibit more static speckle patterns over short time scales (e.g., static regions of fine filipodial extensions). Unlike the remainder of the cell body (which exhibited dynamic speckle patterns), these suppressed structures were not effectively captured by our K-means clustering algorithm. For the purposes of this preliminary study, we assumed that the 3D image segmentation (and resulting 3D meshes) generated by our automated methods provided a sufficiently accurate approximation of the cell surface for quantifying time-varying CTFs. Additional future experiments will be needed to determine whether the loss of fine cellular structures by our algorithm has any significant impact on the accuracy of CTF reconstructions. More advanced algorithms may yield improved cell tracings with higher fidelity^30^, and this aspect of TF-OCM merits further work. Future iterations of TF-OCM may also benefit from the incorporation of hardware-based speckle reduction methods^31,32^. However, the methods presented here do have the advantage that they may be performed using standard OCM microscopes without modification. That is, our methods only require the acquisition of additional images in time, and do not require the introduction of additional hardware or specialized scanning geometries.

### TF-OCM quantifies 3D substrate deformations induced by time-varying CTFs

Unlike the cell body or Matrigel substrate, the polystyrene beads embedded in the sample emitted very strong scattering signals, and so were readily segmented for the purposes of localization and tracking over time. Figure 5 and Supplementary Movie 2 depict the resulting measurements of 3D substrate deformations (in the form of bead displacement data). These data show that, at the beginning of each time-lapse experiment, cell force-induced deformations extended further than 100 μm from the cell body. After the addition of the contractility inhibitor (cytochalasin D) 30 minutes into the experiment, there was a delayed onset of cell relaxation, consistent with our prior work using similar force inhibition protocols^15^. Once relaxation began, bead displacements declined until the beads arrived at their reference (i.e., zero deformation) positions at the end of the experiment. Cell 1 and cell 3 (Figs. 5(a) and 5(c), respectively) demonstrated gradually increasing contractility, until the inhibitor took effect and caused contractility to decline. Cell 2 (Fig. 5(b)), on the other hand, demonstrated a relatively static level of contractility, which then declined in response to the inhibitor. Given the small sample size provided by these three cells, this difference in behavior may be due to multiple sources, such as variable cell health and behavior (e.g. migratory, resting, dividing, etc.). Overall, these results show that TF-OCM offers the capability to measure CTF-induced deformations with minute-scale temporal resolution, enabling the study of dynamic processes, such as the cell relaxation shown here.

### TF-OCM quantifies time-varying, 3D cell traction forces

The results so far have depicted substrate deformation data and cell surface geometry data obtained by our TF-OCM workflow (Fig. 1). After obtaining the mechanical properties of the Matrigel substrate (as described in the Methods), time-varying CTFs were reconstructed using FEM software^26^. Figure 6 depicts CTF reconstructions at a single time point for the three fibroblast cells previously shown in Fig. 5. Animations of these reconstructions are provided in Supplementary Movie 3. Note that cell 1 (shown in Fig. 6(a)) exhibited the greatest degree of polarization, with protrusions extending lengthwise into the surrounding medium. Cell traction forces can be found concentrated at either end of the cell, consistent with prior results from other TFM studies^9,33^. Cells 2 and 3 (shown in Figs. 6(a) and 6(b)) were more spherical in shape. The reconstructed traction fields seem to reflect this relative lack of polarization, as the tractions appear more irregularly distributed across the cell surface, suggesting that a more non-directional contractile behavior is present. Irregular/fluctuating CTF distributions may also result from measurement noise. This is a major challenge for most CTF reconstruction methods, especially in 3D settings^6,9,33^.

Figure 7 summarizes how reconstructed CTFs changed over the course of each time lapse experiment. Each data point represents the total force exerted by a cell at a given time point, oriented along its principal axis of stress (detailed in the Supplementary Methods). These plots show that the forces exerted by the fibroblast cells tended to increase until approximately 30 minutes after the addition of the contractility inhibitor, after which their contractility declined. These curves also resemble the deformation distributions in Fig. 5, as would be expected. The variability between individual time points (both in Supplementary Movie 3 and Figure 7) may be due to a combination of factors, including actual changes in CTF distribution, noise, and artifacts emerging from the use of FEM. Overall, these data have demonstrated the ability of TF-OCM to measure trends in time-varying CTFs exerted by cells with minute-scale temporal sampling.

## Discussion

As shown by the results in Fig. 2 and Supplementary Movie 1, the phase registration and bulk demodulation techniques used in this study mitigate distortions that can appear in volumetric images reconstructed with CAO. A detailed discussion of the origin of these distortions may be found in the Supplementary Discussion. In brief, the shearing artifact shown in Fig. 2 originates from a tilt/misalignment in the optics and/or sample stage, while the motion artifacts visible in Supplementary Movie 1 originate from interferometric phase instability. The effects of tilt must be mitigated either through signal demodulation (as was done in this study) or a modification of the equations underlying CAO (specifically, Eqn. (S.10) in the Supplementary Methods). The effects of phase instability require special attention from any who wish to perform TF-OCM. It is well known that CAO and related computed imaging methods rely on the use of phase stable OCM image data^23,24^. The discussions surrounding phase stability typically focus on the ability of CAO to reconstruct high resolution volumetric images. That is, phase instability may be regarded as a negligible factor to image formation if high resolution images can be restored. However, our results show that applications like TF-OCM have stricter phase stability requirements than applications which solely require high spatial resolution. This is because, although a small degree of phase instability may not noticeably impact the final spatial resolution, it can still result in image distortion artifacts like those seen in Supplementary Movie 1, which are detrimental to TF-OCM and other applications which use reconstructed images to measure features of the sample structure. Therefore, phase stabilization is an essential component of the TF-OCM processing workflow, even for imaging systems which exhibit a high degree of phase stability.

TFM relies on the ability to perform accurate quantitative analysis of substrate deformations. The implementation of TF-OCM in this work relied upon a number of coordinate transformations (in both the space and spatial frequency domains) to achieve its objectives. These operations (shown in Fig. 1, and detailed in the Supplementary Methods) included standard OCM image reconstruction procedures, coherence gate curvature removal, focal plane registration, phase registration, bulk demodulation, CAO, and motion correction. Each of the steps in our computational image formation module involved coordinate transformations and/or phase manipulations which can have dramatic effects on the final image structure, substrate deformation data, cell tracings, and CTF reconstructions. As a result, these steps offer several opportunities for errors to emerge before the final image structure is generated. Optical distortions and the question of image fidelity are problems faced when using any type of imaging system. Whether formed physically or through computation, an image can only be a best attempt to represent the true underlying sample structure. To provide further insight into the workings, reasoning, and potential flaws of our TF-OCM methods, major assumptions and possible failure modes of our processing workflow have been detailed in the Supplementary Discussion.

One potential disadvantage of the experimental methods shown here is the use of relatively large (1 μm diameter) scattering beads. High concentrations of beads allow for substrate deformations to be sampled more densely in space, and therefore can help to mitigate the impact of error/measurement noise. In general, beads used for TFM are typically at least one order of magnitude smaller than the minimum desired bead spacing^19,33,34^. Using 1 μm beads allows for a minimum bead spacing of approximately 10 μm (the actual spacing in the experiments reported here was approximately 18 μm). Future experiments with TF-OCM may use smaller beads to allow for greater bead densities without disrupting substrate mechanics or cell behavior. However, as a decrease in bead size tends to decrease the strength of the scattered signal from a given bead, any reduction in bead size must optimize a trade-off between bead density and SNR of the bead signal with respect to the imaging noise floor. In addition, the presence of highly scattering particles may obstruct any co-registered fluorescence imaging which may accompany a TF-OCM experiment. One solution to this problem may be to abandon scattering beads entirely. TFM has been demonstrated using confocal reflectance microscopy to track the motion of collagen fibers, instead of embedded beads^35^. OCT/OCM could similarly be used to image the deformation of fibrous extracellular matrix constituents for TF-OCM. If the scattering structures (e.g., collagen fibers) are too small to resolve, speckle tracking methods (analogous to those used for optical coherence elastography^36^), may be a possible solution.

Speckle is another limitation of TF-OCM. This study required the use of speckle reduction methods to perform TF-OCM based on automated analysis of time-lapse OCM image data. Future implementations of TF-OCM based purely upon OCT/OCM imaging will likely continue to rely on speckle reduction procedures^30-32^. However, TF-OCM could also be performed in conjunction with co-registered fluorescence imaging to capture images of cell geometry without speckle. If the cell shape changes slowly in time, fluorescence imaging would not have to take place at every time point, so as not to limit the temporal resolution provided by the rapid volumetric acquisition of OCT/OCM imaging. Moreover, fluorescence imaging may be confined solely to the cell/s of interest while OCM is used to capture the full volumetric FOV under study, thereby keeping any photobleaching/phototoxicity from fluorescence imaging to a minimum during extended experiments.

Similar to most TFM procedures, the reconstruction of CTFs in this study relied on several assumptions about the substrate medium (e.g., that the material is linear elastic, and its mechanical properties are isotropic, homogeneous, and time-invariant). In practice, biopolymer substrates often violate these assumptions. Nonlinear materials can be accommodated through the use of alternative mechanical models^19^. However, measurements of any spatial or temporal variations in local mechanical properties near cell bodies are difficult to obtain in 3D settings. TF-OCM is readily compatible with optical coherence elastography (OCE)^36^, as both methods share OCT as a parent imaging modality. In the future, TF-OCM may benefit from integration with high resolution OCE methods that can measure local variations in substrate mechanical properties^37,38^.

## Conclusion

In this study, we have implemented quantitative TF-OCM for the first time, enabling the reconstruction of time-varying 3D cell traction forces with minute-scale temporal resolution. This was achieved by combining rapid OCM image acquisition with computational image formation methods, applied to an otherwise standard TFM experimental protocol. As a result, TF-OCM has been demonstrated to be a viable method for overcoming the limitations of typical TFM methods by providing a large volumetric coverage, high temporal resolution, and a low risk of photobleahing/phototoxicity. Due to the reliance of high-throughput 3D TF-OCM on computational methods, we provided extensive discussion of these methods, their assumptions, and potential drawbacks. Currently, TF-OCM has only been applied to single cells. However, due to the ability of OCT/OCM to image over large volumes in scattering media, TF-OCM may find its most impactful niche in the study of physiologically-relevant multicellular constructs on spatial scales ranging from hundreds of micrometers to millimeters, such as cell networks and spheroids. Since OCM is capable of imaging native scattering contrast, future implementations of TF-OCM may also enable the study of CTFs in systems that do not have fiducial marker beads, a capability that is currently uncommon in the TFM field. Moreover, TF-OCM is compatible with optical coherence elastography, making OCT imaging systems an attractive platform for mechanobiology research as both techniques continue to develop. Finally, due to the highly modular nature of the TF-OCM protocol, the procedures outlined in this report may be readily improved, modified, or adapted to a variety of alternative settings and imaging preferences. Consequently, TF-OCM has significant potential for making contributions to research in mechanobiology.

## Methods

### Sample preparation

Samples consisted of NIH-3T3 fibroblasts embedded in Matrigel containing scattering polystyrene beads. Cells were maintained in tissue culture flasks with media consisting of Dulbecco’s Modified Eagle Medium (DMEM), supplemented with 10% bovine calf serum, and 1% penicillin-streptomycin. To prepare samples, the cells were trypsinized, pelleted, and resuspended in chilled (4 °C) media at a concentration of 3.33×10^4^ cells/mL. 1 μm-diameter polystyrene beads (Spherotech Inc.) were added to the suspension to achieve a concentration of 5.3×10^8^ beads/mL. The cell-bead suspension was added to Matrigel in a 30:70 ratio, resulting in a final cell concentration of 1×10^4^ cells/mL, and final bead concentration of 1.6×10^8^ beads/mL (yielding an average bead spacing of approximately 18 μm). The resulting mixture was deposited in 100 μL aliquots on glass-bottomed petri dishes and left to gel in an incubator for 20 minutes, before being covered in culture media. Samples were kept in an incubator overnight (~12 hours) prior to imaging.

### Bulk rheology

Sample stiffness was characterized using a TA Instruments DHR3 shear rheometer in a (flat) plate-plate geometry. Cell culture media and Matrigel were mixed in a 30:70 ratio. For each test, the mixture was loaded and gelled underneath a 40 mm diameter plate on a temperature controlled testing stage at 37 °C for 20 minutes. The sample stage was covered/sealed to mitigate evaporation. Approximating the Matrigel hydrogel as a nearly incompressible substance (with Poisson’s ratio=0.45), the Young’s modulus was found to be approximately 90 Pa. This value was used as an input to a Finite Element Method solver to reconstruct CTFs.

### Imaging system

All samples were imaged using a spectral domain OCM (SD-OCM) system. Illumination was supplied by a Ti:Sapph laser (Femtolasers, INTEGRAL Element) with a central wavelength of 800 nm and a bandwidth of 160 nm. Light was split between the sample and reference arms by a 90:10 fiber coupler, yielding an incident power of ~5 mW in the sample arm beam. The sample arm was built in a double-pass configuration with an Olympus XLUMPlanFL 20×/0.95W ∞/0 objective lens in an inverted configuration (i.e., samples were imaged through the bottom of the glass-bottomed petri dish). Spectral data was acquired with a Cobra 800 spectrometer (Wasatch Photonics) and 2048-pixel line scan camera (e2v, Octopus). The axial and transverse resolutions of the system were approximately 2.4 μm and 1.5 μm, respectively. All data acquisition was performed with a line scan rate of 65 kHz and an exposure time of 10 μs. Under these conditions, the system exhibited a sensitivity of ~90 dB and fall-off of −5 dB/mm. The system was controlled using custom software built in LabVIEW.

### Time-lapse imaging protocol

Samples were held in position by an Okolab UNO-PLUS incubating stage mounted on a (non-motorized) 3-axis translation stage. This incubating stage maintained physiological temperature, humidity, and pH levels in the samples during imaging. Due to the use of a non-motorized stage, only a single embedded fibroblast cell was imaged per time-lapse experiment. For each time-lapse, an isolated cell (i.e., a cell which occupied its own ‘personal’ 500×500×500 μm^3^ FOV within the Matrigel substrate) was located. This cell was aligned to the focal plane of the OCM system and centered within the transverse FOV. Time-lapse imaging was then begun. Each time-lapse imaging experiment spanned a total of 3 hours. This 3 hour period was divided into two phases. The first phase spanned the first 30 minutes of imaging. During this time, the baseline (contractile) behavior of the cell was monitored. At the end of this phase, a contractility inhibitor (0.5 mM cytochalasin D dissolved in DMSO) was added to the petri dish to achieve a final cytochalasin D concentration of 1 μM. The second phase spanned the remaining time of the experiment (2.5 hours). During this phase, cell relaxation in response to the contractility inhibitor was observed.

Bursts of volumetric images were acquired at five-minute intervals across both phases of each time-lapse experiment. Each burst captured a single ‘full FOV’ volume (spanning 2560×500×500 μm^3^ with 2048×1024×1024 voxels) and eight ‘reduced FOV’ volumes (spanning 2560×125×125 μm^3^ with 2048×256×256 voxels). This multi-acquisition/multi-FOV scheme was used to aid in speckle reduction for automated segmentation of cell bodies. Each burst of images took approximately 1.5 minutes to acquire. The laser shutter was closed between each burst to limit laser exposure of the sample between acquisitions.

### Computational image formation

The reconstruction of volumetric time-lapse images from raw OCM spectral data consisted of six steps: 1) initial volume reconstruction, 2) coherence gate curvature removal, 3) focal plane registration, 4) phase registration, 5) bulk demodulation, and 6) computational adaptive optics. This procedure was designed to provide accurate high resolution volumetric image data that is well-suited for quantitative TFM applications. For a given time-lapse dataset, our implementation of this procedure processed each time point in series. However, it should be noted that this procedure is also amenable to parallelization. All steps were performed in MATLAB R2014b using CPU-based processing. In addition, all procedures were automated, requiring no human input except where otherwise stated in the complete method descriptions, which are available in the Supplementary Methods.

### Measurement of substrate deformations

Substrate deformations were determined from the motion of the scattering polystyrene beads embedded in the Matrigel substrate. Bead motion was tracked using a time series consisting of the ‘full FOV’ images (described previously) in order to capture substrate deformations both near and far away from the cell surface.

As the scattering polystyrene beads exhibited a high signal strength well above the noise floor, bead positions were localized with a simple segmentation procedure. Volumetric image intensities were first normalized across depth. This was followed by binarization via single-level thresholding. The binarized images were then cleaned to remove objects that were too small or too large to be scattering beads. Objects removed included those smaller than 16 voxels (i.e., those too small to be a bead), and those larger than the 99^th^ percentile of all objects larger than 16 voxels (such as the cell body, bead aggregates, or protein debris). All remaining objects in the binarized images were assumed to be beads. The location of each bead, was measured by calculating the intensity-weighted centroid of each object in the binary image. The sensitivity of this bead localization procedure was determined by measuring the standard deviation of apparent (i.e., measured) bead displacements between two sequential images, acquired in the absence of CTFs. The localization sensitivity was found to be 86 nm, 32 nm, and 37 nm along the vertical, fast, and slow axes of the imaging system, respectively. (These values correspond to approximately one-fifteenth of the voxel size along each dimension.)

Bead motion over time was determined using a feature vector-based point tracking algorithm^39^. In brief, the algorithm searches for pairs of bead positions (spanning pairs of time points) which are most likely to correspond to the same bead. Candidate pairs of positions are then determined to be a ‘match’ based upon the relative motion of other nearby beads (i.e., the algorithm assumes the substrate behaves as an elastic solid with deformations varying on scales longer than the bead spacing). Using this algorithm, bead positions were tracked across the full temporal span of each time-lapse dataset.

Once the positions of all individual beads were determined across time, bead displacements were calculated with respect to the final bead positions (i.e., the reference positions, as discussed in the ‘Overview of TF-OCM’ section) at the end of the experiment (i.e., time *t* = *t*_max_, or *t* = 3 hours, in our imaging procedure). Details about this procedure, as well as motion correction, may be found in the Supplementary Methods. We assumed both that the fibroblast cells exerted no forces and that the substrate had no internal strain remaining at the end of each time-lapse experiment. This is a standard practice in TFM experiments which use chemical reagents to inhibit cell contractility or induce cell death^9^.

### Image segmentation for the measurement of time-varying cell geometry

Knowledge of the time-varying cell shape and location was required in order to provide the boundary conditions necessary for CTF reconstruction. The 8 ‘reduced FOV’ images (described previously) at each time point were used to reduce speckle, and thereby enable a simple automated image segmentation procedure to identify the cell body. Due to the motion of intracellular components, the speckle pattern varied between individual images. At each time point, the corresponding set of reduced FOV images was combined via a projection operation. Specifically, the (speckle-reduced) output image was obtained as the standard deviation of the magnitude of the corresponding reduced FOV images, taken on a voxel-by-voxel basis. This caused regions of high speckle fluctuation to be emphasized, and regions of static background scattering to be suppressed. It also had the effect of reducing the speckle contrast within the cell body. K-means clustering of a given speckle-reduced volumetric image yielded a binary image from which a volumetric approximation of the cell body was obtained (see Fig. 4).

### Mesh generation

For each time point, the binary image generated by image segmentation was converted into a discretized 3D triangular surface mesh using the *iso2mesh*^40^ and ‘Smooth Triangulated Mesh’^41^ packages in MATLAB (see the Supplementary Methods for details about motion correction). This surface mesh was then used to generate a volumetric tetrahedral mesh of the imaged volume in the open source program, Gmsh^42^. The outer surface of the mesh was a cube defined by the image boundaries, and the inner surface was defined by the cell surface at that time point. Finally, this volumetric tetrahedral mesh was converted into a volumetric hexahedral mesh using the open source software package, *tethex*^43^. The final output of this procedure was a sequence of volumetric meshes suitable for use in our CTF reconstruction software.

### Reconstruction of time-varying 3D cell traction forces

The substrate mechanical characterization data, bead displacement data, and 3D mesh data produced by all the methods described above were used to reconstruct time-varying 3D cell traction forces. CTF reconstruction was performed using a previously reported custom FEM software package^26^ based on the open-source deal. II library^44^. In brief, the reconstruction of cell traction forces poses an inverse problem, which was solved by minimizing the discrepancy between the measured bead displacements and those that would be predicted to result from a candidate hypothesized traction field. The Matrigel substrate was assumed to be linear elastic, homogeneous, isotropic, and time-invariant. (These are standard, although neither universal nor mandatory, assumptions used to make the inverse problem posed by traction force reconstruction tractable^9^.) As cell motion was quasi-static for any given time point, the reconstructed traction field was required to satisfy force and moment balance. To optimize the trade-off between accuracy and instability of the numerical solution, Tikhonov regularization was applied (using a regularization coefficient value of 10^−7^, which was determined using the L-curve method)^26^. The mechanical properties used as input were a Young’s modulus of 90 Pa (measured, as described previously) and Poisson’s ratio of 0.45 (assumed). CTF reconstruction was carried out independently for each time point in a given time-lapse experiment.

## Data availability

All relevant data are available from the authors.

## Acknowledgements

The authors would like to thank Nichaluk Leartprapun for performing bulk rheology tests, and Dr. Warren Zipfel for loaning the objective lens used for this study. This research was supported in part by a grant from the National Institutes of Health (NIBIB-R21EB022927) awarded to SGA. This study made use of the Cornell Center for Materials Research Shared Facilities, which are supported through the NSF MRSEC program (DMR-1120296).

## Author contributions

J.A.M. wrote/generated all materials (text, supplementary information, figures, and movies) of this report. J.A.M. performed all cell culture, sample preparation, image data acquisition, and data processing. J.A.M. developed all components of the computational image formation module. J.A.M. developed the bead localization, motion correction, speckle reduction, and image segmentation components of the traction force microscopy module. X.F. devised the algorithm for bead tracking, and J.A.M. developed a MATLAB implementation of the algorithm. X.F. developed the mesh generation and force reconstruction components of the traction force microscopy module. S.G.A. provided guidance and funding throughout this study. All authors reviewed and edited the full contents of this report.

